# Nascent CUT&Tag captures transcription factor binding after chromatin duplication

**DOI:** 10.1101/2025.10.13.682212

**Authors:** Matthew Wooten, Kevin Nguyen, Brittany N. Takushi, Kami Ahmad, Steven Henikoff

## Abstract

DNA replication strips off all chromatin proteins, which must be reassembled behind the replication fork. To track chromatin reassembly on newly synthesized DNA, we developed Nascent CUT&Tag, a chromatin profiling method that directly measures transcription factor (TF) binding on nascent chromatin. We tracked the recovery of GAGA factor (GAF) and Pleiohomeotic (PHO) in Drosophila Kc167 cells. Whereas both TFs are evicted following passage of the replication fork, GAF recovers on newly synthesized DNA over a minutes-to-hours range, whereas PHO generally requires hours to fully reestablish binding. Early recovering GAF peaks are characterized by shorter GAF motifs and are associated with functions related to cell cycle progression. Conversely, late recovering peaks are characterized by longer, degenerate GAF motifs and are associated with developmental functions. GAF recovery on newly synthesized DNA requires chromatin remodeling by Brahma Associated Factor (BAF), implying that nucleosome turnover is critical to fully reestablish GAF binding.

## Introduction

The majority of DNA in eukaryotic genomes is organized into nucleosomes, which are comprised of 146 base pairs of DNA wrapped around the histone octamer^1^. Transcription factors (TFs) are DNA binding proteins that alter chromatin structure and regulate gene expression by binding to targeted DNA sequences^2,3^. Nucleosome occupancy antagonizes TF binding, as most TFs cannot recognize and bind the target motifs wrapped around nucleosomes. DNA replication poses a formidable challenge to TF binding, as TFs become displaced from DNA following passage of the replication fork and replaced by nucleosomes^4,5^. To reestablish parental chromatin structure, TFs must overcome nucleosome occlusion to access and bind to their target motifs. However, despite the critical role that TFs play in regulating chromatin structure and gene expression, little is known regarding the kinetics or mechanisms by which TFs reassociate with DNA following passage of the replication fork.

To directly track the return of chromatin features to newly synthesized DNA, we developed a chromatin profiling method called Nascent CUT&Tag, which uses targeted *in* situ tagmentation to directly measure transcription factor binding on EdU-labeled nascent chromatin^6^. We used Nascent CUT&Tag to track the return of GAGA factor (GAF) and Pleiohomeotic (PHO) in *Drosophila melanogaster* Kc167 cells. GAF is an essential *Drosophila* zinc finger TF encoded by the Trithorax-like (*Trl*) gene^7^ that binds to GA-rich sequence arrays on the promoters and cis-regulatory elements^8^ to drive gene activation and repression^9,10^. PHO is a zinc finger transcription factor that binds to DNA and interacts with members of the Polycomb repressive complex to help establish and maintain gene repression^11–13^. Using Nascent CUT&Tag, we find that GAF and PHO are displaced from chromatin during DNA replication and recover binding over time. PHO shows delayed recovery kinetics, with most PHO sites requiring hours to fully reestablish PHO occupancy. GAF shows a range of recovery kinetics on newly synthesized DNA, with certain sites recovering binding minutes after fork passage, while other sites recover binding hours later. Early recovering GAF peaks contain short GAF DNA motifs and are located at genomic features associated with cell cycle progression. Conversely, late recovering GAF sites are characterized by longer, more degenerate GAF motifs and are found at genomic features associated with developmental functions. Using the small molecule inhibitor BRM014, we show that BAF activity is essential for GAF to fully recover binding to nascent chromatin. Together, this work reveals a role for nucleosome remodeling in facilitating TF binding to newly synthesized DNA and suggests that motif structure plays a role in GAF recovery on nascent chromatin.

## Results

### Nascent CUT&Tag profiles chromatin features on newly synthesized DNA

To detect factor binding on nascent DNA, we combined pulse-labeling of newly synthesized DNA with Cleavage Under Targets and Tagmentation (CUT&Tag) chromatin profiling (**Figure 1A).** We pulse-label Kc167 cells with a 15-minute treatment of 5-Ethynyl-2’-deoxyuridine (EdU), followed by increasing chase times to capture factor binding at different stages of chromatin maturation. Cells are then lightly fixed and processed for CUT&Tag with antibodies to a chromatin factor. Following tagmentation, we use click chemistry to link a biotin moiety to incorporated EdU, and then streptavidin pulldown to purify replicated DNA. PCR with tagmentation primers thus generates sequencing libraries specifically from replicated chromatin. Libraries were pooled by equal volumes to accurately represent amounts of chromatin-bound factors between timepoints.

**Figure 1.**
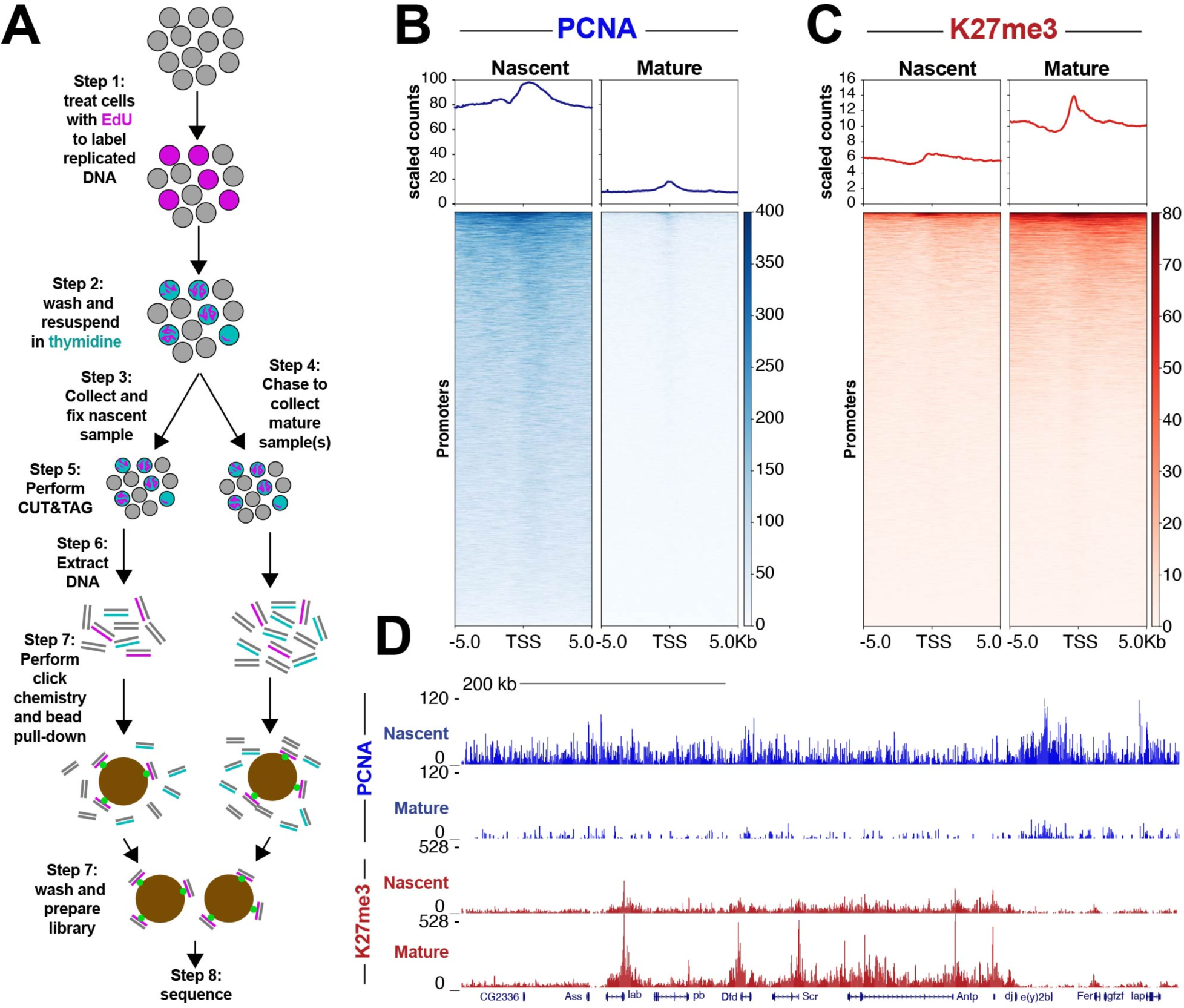
Nascent CUT&Tag captures changes in PCNA and H3K27me3 behind the replication fork. (**A**) Experimental design for Nascent CUT&Tag workflow. (**B**) Heatmap aligned to transcription start site (TSS) of all promoters showing PCNA counts scaled by number of mapped reads. (norm.). (**C**) Heatmap aligned to transcription start site (TSS) of all promoters showing H3K27me3 counts scaled by number of mapped reads. (**D**) Representative UCSC browser tracks snapshot of PCNA and H3K27me3. Merged data of three biological replicates for PCNA and four biological replicates for H3K27me3.

To validate Nascent CUT&Tag, we first tested if we could detect the replication-coupled processivity factor PCNA on new DNA, as this should be specifically enriched in nascent libraries^14,15^. Indeed, we observe strong PCNA signal on newly synthesized DNA, while more mature chromatin shows reduced signal (**Figure 1B,D**). To further validate Nascent CUT&Tag, we measured the maturation kinetics for trimethylation at the histone H3 K27 residue (H3K27me3). This histone modification is catalyzed by the Polycomb repressive complex (PRC2) and promotes gene repression. Levels of H3K27me3 are initially diluted by half following passage of the replication fork, as duplicated chromatin contains an approximately equal mixture of old histones with the H3K27me3 mark and new histones that are unmarked. Over the course of hours, PRC2 methylates new histones, leading to a 2-fold increase in K27me3 levels from nascent chromatin to mature^16,17^. Indeed, H3K27me3-targeted Nascent CUT&Tag signals are low on newly replicated chromatin and then increase approximately two-fold on mature chromatin (**Figure 1C,D**). These experiments demonstrate that Nascent CUT&Tag quantitatively profiles diverse chromatin features on newly synthesized DNA.

### GAF recovers rapidly on newly synthesized DNA

Nascent chromatin must mature to reestablish nucleosome organization and transcription factor binding. To describe the kinetics, we chose to profile the *Drosophila* transcription factors PHO and GAF. GAF is an abundant zinc-finger transcription factor that is crucial for developmental gene expression programs in *Drosophila*. GAF binds at thousands of gene promoters, developmental enhancers, as well as heterochromatic satellite blocks that have AAGAG and other GA-rich motifs^10,18^. From bulk profiling, we annotated ∼7,000 sites throughout the genome of *Drosophila* Kc167 cells (**Supplementary Table 1**). To then assess GAF binding to nascent DNA, we pulsed-labeled cells with EdU and then immediately prepared cells for profiling (T0) or chased cultures with thymidine for 1 or 4 hours (T1 and T4). These timepoints capture newly synthesized chromatin and progressively more mature chromatin after replication.

Overall, 50% of GAF signal is rapidly restored on newly synthesized DNA and the remainder gradually recovers over 4 hours (**Figure 2A**). However, at some sites recovery varies: We identified 265 sites that more rapidly acquire GAF signal and 157 sites that recover more slowly (**Figure 2B**). The 265 early-recovery sites have relatively low signal, but these sites appear maximally occupied immediately after replication fork passage (**Figure 2C,E**). By contrast, the late-recovery sites are more prominent GAF peaks, but have very low signal on new chromatin, and gradually increase to their maximum after 4 hours (**Figure 2D,G**). While early recovering peaks have a short GAF consensus motif (**Figure 2F**), late recovering sites contain longer, more degenerate GAF motifs (**Figure 2H**), suggesting that motif structure plays a role in the timing of GAF recovery. Early and late recovery peaks also show a substantial difference in peak width, with late recovery peaks showing a median peak width of 1603 basepairs (bp), while early recovery peaks show a median peak width of 942 bp (**Figure 2C** *versus* **Figure 2D**).

**Figure 2.**
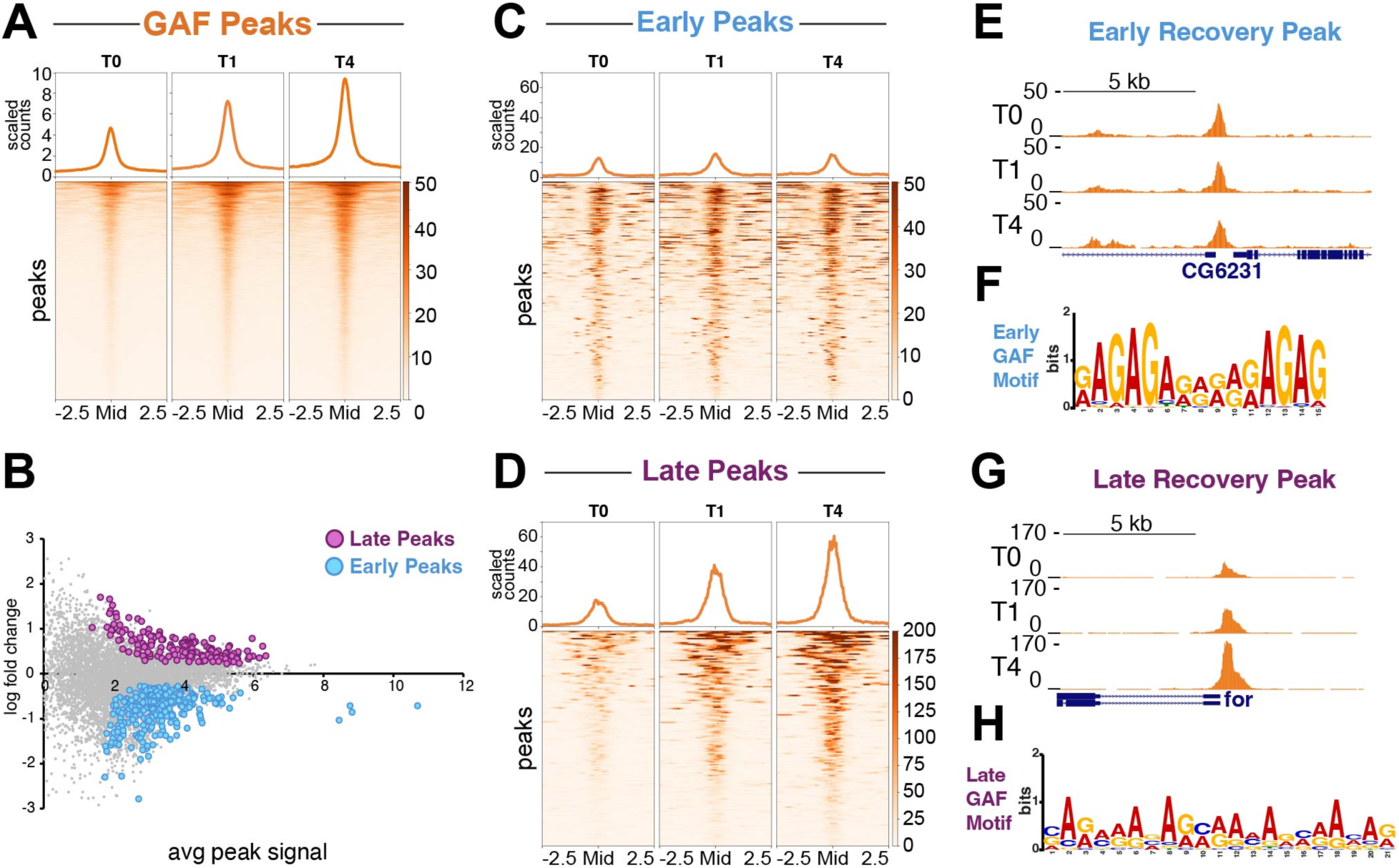
GAF binding is lost on nascent chromatin and shows early and late recovery. (**A**) Heatmap aligned to the center of all GAF peaks showing GAF signal over the course of Nascent CUT&Tag experiment. T0 is newly synthesized chromatin immediately after replication fork passage. T1 is one hour after replication fork passage and T4 is four hours after replication fork passage. Read counts are scaled by number of mapped reads. (**B**) MA plot showing fold change in normalized counts comparing T4 to T0. Statistical significance determined by two-sided t-test. (**C**) Heatmap aligned to the center of all early recovering GAF peaks showing GAF signal over the course of Nascent CUT&Tag experiment (**D**) Heatmap aligned to the center of late recovering GAF peaks showing GAF signal over the course of Nascent CUT&Tag experiment. (**E**) Representative UCSC browser track snapshot centered on the promoter of CG6231 showing GAF signal across Nascent CUT&Tag time course for early recovering peak. (**F**) MEME suite motif analysis of early recovering peaks. (**G**) Representative UCSC browser track snapshot centered on the promoter of *foraging* showing GAF signal across Nascent CUT&Tag time course for late recovering peak. (**H**) MEME suite motif analysis of late recovering peaks. Merged data of ten biological replicates for T0, six biological replicates for T1 and seven biological replicates for T4.

We also performed differential analysis on *Drosophila melanogaster* features comprised of promoters, enhancers, and Polycomb response elements (PREs), which are regulatory DNA elements that bring Polycomb group proteins to DNA. We identified 51 (38%) promoters, 41 (30%) house-keeping enhancers, 35 (26%) developmental enhancers and 8 (6%) PREs that showed early recovering kinetics. We also identified 167 (49%) promoters, 74 (21%) house-keeping enhancers, 97 (29%) developmental enhancers and (<1%) 2 PREs (**Supplementary Figure 1**) that showed later recovering kinetics. Early-recovery promoters are enriched for biological function related to the cell cycle, while late-recovery promoters were associated with developmental functions (**Table 1**). This suggests that slow recovery may be a feature important for regulation of genes active in distinct cell types.

**Table 1:**
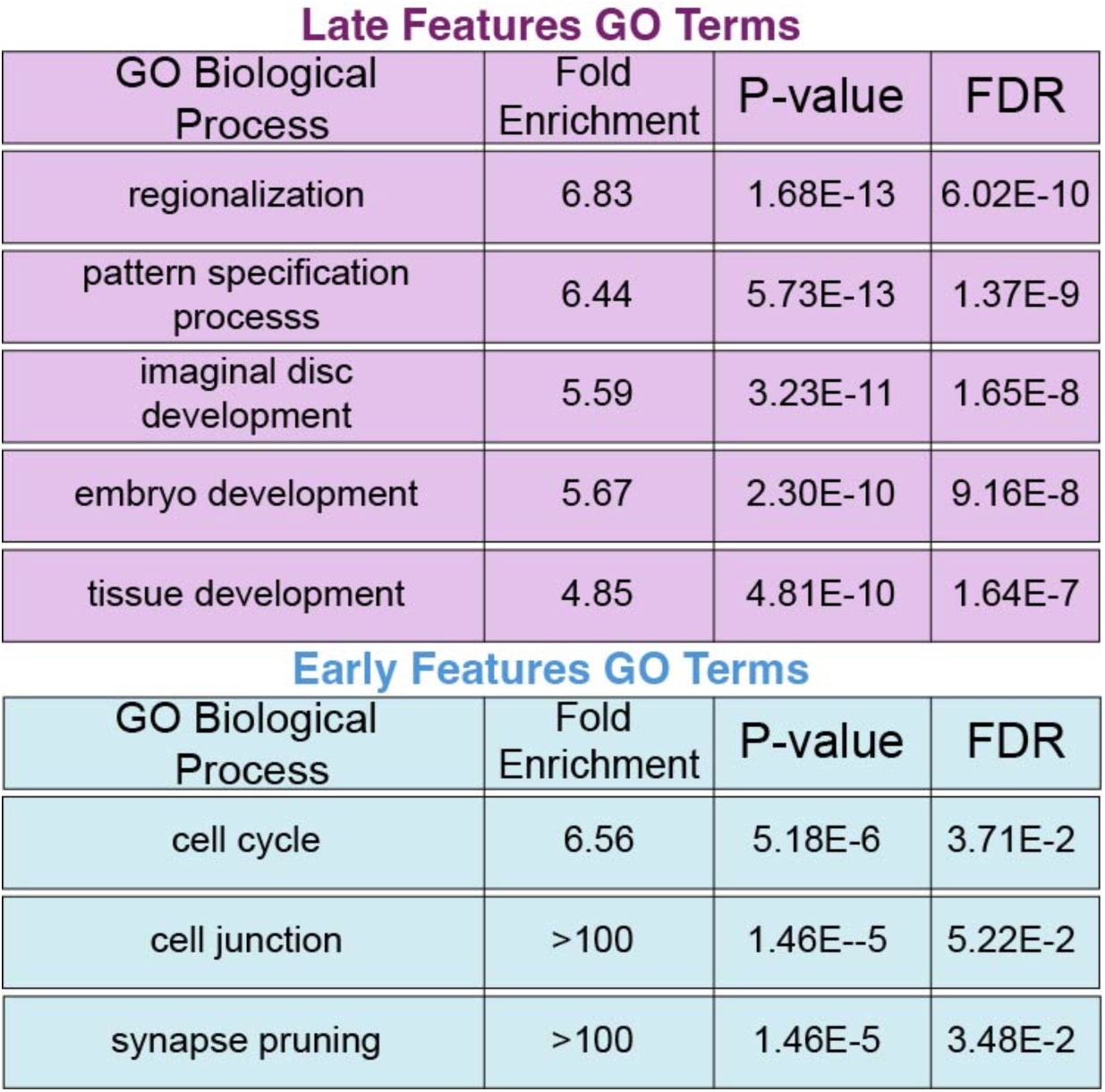
Gene Ontology (GO) term analysis of early and late recovering GAF features.

Late Features GO Terms
| GO Biological Process | Fold Enrichment | P-value | FDR |
| --- | --- | --- | --- |
| regionalization | 6.83 | 1.68E-13 | 6.02E-10 |
| pattern specification processs | 6.44 | 5.73E-13 | 1.37E-9 |
| imaginal disc development | 5.59 | 3.23E-11 | 1.65E-8 |
| embryo development | 5.67 | 2.30E-10 | 9.16E-8 |
| tissue development | 4.85 | 4.81E-10 | 1.64E-7 |

| GO Biological Process | Fold Enrichment | P-value | FDR |
| --- | --- | --- | --- |
| cell cycle | 6.56 | 5.18E-6 | 3.71E-2 |
| cell junction | >100 | 1.46E--5 | 5.22E-2 |
| synapse pruning | >100 | 1.46E-5 | 3.48E-2 |

### PHO recovers slowly on newly synthesized DNA

PHO is a zinc finger transcription factor involved in gene silencing that interacts with Polycomb proteins and binds to PREs. From bulk profiling, we annotated ∼3,000 sites of PHO binding throughout the genome of *Drosophila* Kc167 cells (**Supplementary Table 2**). To then assess PHO binding to nascent DNA, we pulsed-labeled cells with EdU and then immediately prepared cells for profiling (T0) or chased cultures with thymidine for 1 or 4 hours (T1 and T4).

In contrast to GAF where 50% of GAF occupancy is rapidly restored, PHO showed minimal recovery at its sites immediately after replication fork passage, as newly replicated DNA (T0) shows only 30% of the global occupancy measured after 4 hours (T4) maturation (**Figure 3A**). PHO undergoes limited recovery even after 1hr of chromatin maturation, as the T1 timepoint still shows only 44% of the global PHO occupancy detectable at T4, indicating that that majority of PHO recovery occurs hours after replication fork passage. We further identified 245 peaks that showed significant delays in PHO binding to nascent chromatin (**Figure 3B**). PHO binding at these sites was barely detectable immediately (13% recovery) after the replication fork (T0), indicating the PHO is unable to regain occupancy on newly synthesized DNA. PHO binding at these sites showed minimal gains in occupancy after one hour of chromatin maturation (23% recovery), indicating that PHO recovery at these sites is substantially delayed (**Figure 3C,D**). To directly compare PHO and GAF recovery at late recovering PHO sites, we plotted GAF occupancy at late recovering PHO sites. In contrast to PHO, we observed substantial GAF occupancy at these sites immediately after replication fork passage (45% recovery) and 1hr later (73% recovery), indicating that GAF reestablishes binding at PHO late recovery sites more rapidly than PHO (**Figure 3E, F)**.

**Figure 3.**
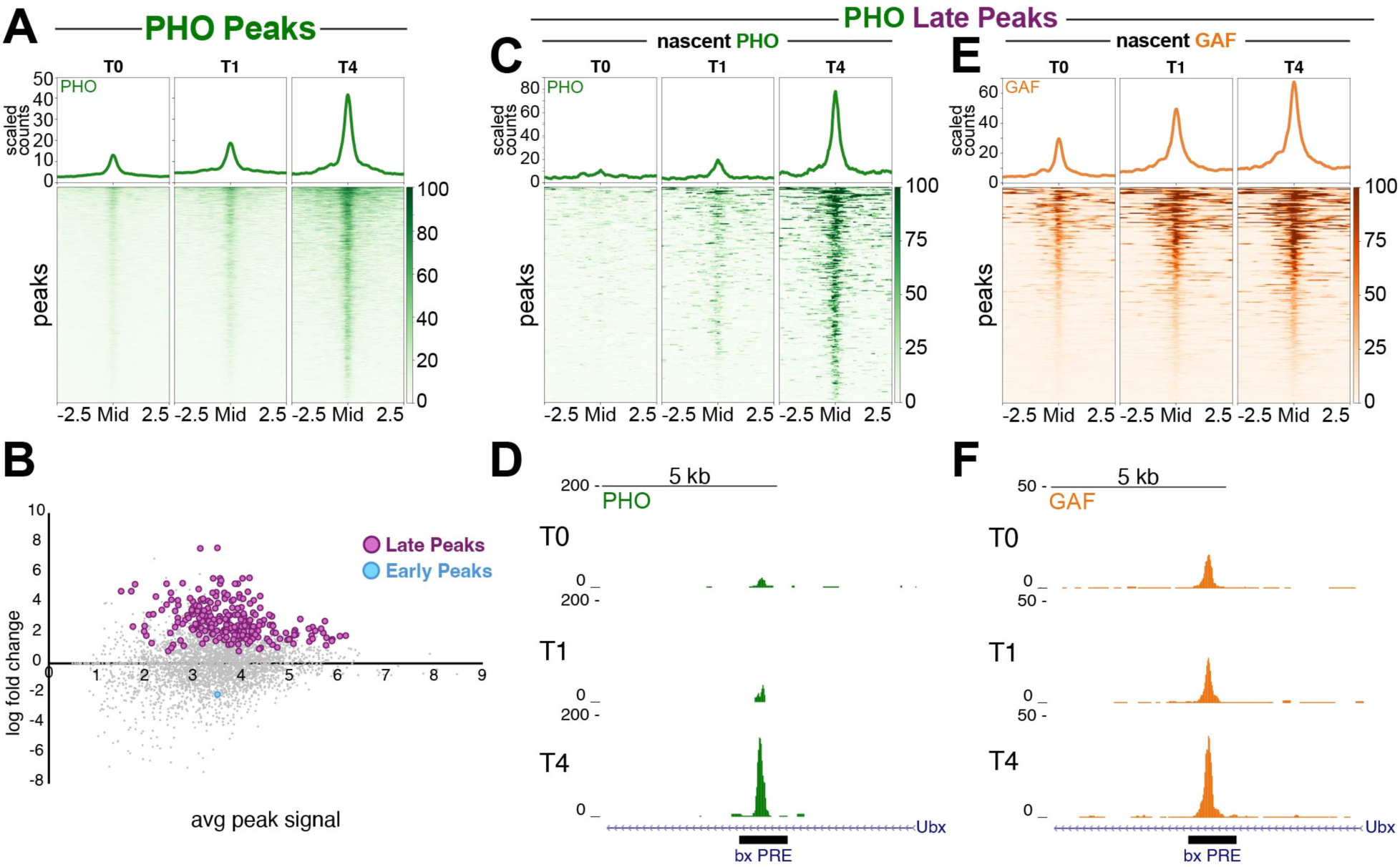
PHO binding is lost on nascent chromatin and shows late recovery. (**A**) Heatmap aligned to the center of all PHO peaks showing PHO signal over the course of Nascent CUT&Tag experiment. T0 is newly synthesized chromatin immediately after replication fork passage. T1 is one hour after replication fork passage and T4 is four hours after replication fork passage. Read counts are scaled by number of mapped reads. (**B**) MA plot showing fold change in normalized counts comparing T4 to T0. Statistical significance determined by two-sided t-test. (**C**) Heatmap aligned to the center of late recovering PHO peaks showing PHO signal over the course of Nascent CUT&Tag experiment. (**D**) Representative UCSC browser track showing PHO coverage centered on the *bx* Polycomb Response Element (PRE) in the *Ultrabithorax* (*Ubx*) gene. (**E**) Heatmap aligned to the center of late recovering PHO peaks showing GAF signal over the course of Nascent CUT&Tag experiment. (**F**) Representative UCSC browser track showing GAF coverage centered on the *bx* Polycomb Response Element (PRE) in the *Ultrabithorax* (*Ubx*) gene. Merged data of six biological replicates for T0, three biological replicates for T1 and three biological replicates for T4.

### Chromatin remodeling is required for transcription factor recovery on nascent chromatin

It is surprising that at some sites transcription factor binding is delayed by hours after passage of the replication fork. Previous studies have suggested that nucleosomes behind the fork may occlude factor binding sites^4,5^. We wondered if chromatin must be remodeled at some genomic sites behind the replication fork to expose factor motifs, and this might delay factor binding. This model implies that chromatin remodeling may be important for chromatin maturation.

Specifically, the BAF chromatin remodeler is enriched at promoters that transiently gain nucleosomes that are later repositioned^5^. To assess if BAF is similarly involved at GAF binding sites, we profiled the Moira (BAFp170 homolog) subunit of BAF complexes^19^ using a modified high-yield CUT&Tag protocol^20^. We then plotted Moira coverage at all GAF sites and observed clear Moira enrichment, indicating that BAF associates with GAF-bound sites (**Figure 4A**).

**Figure 4:**
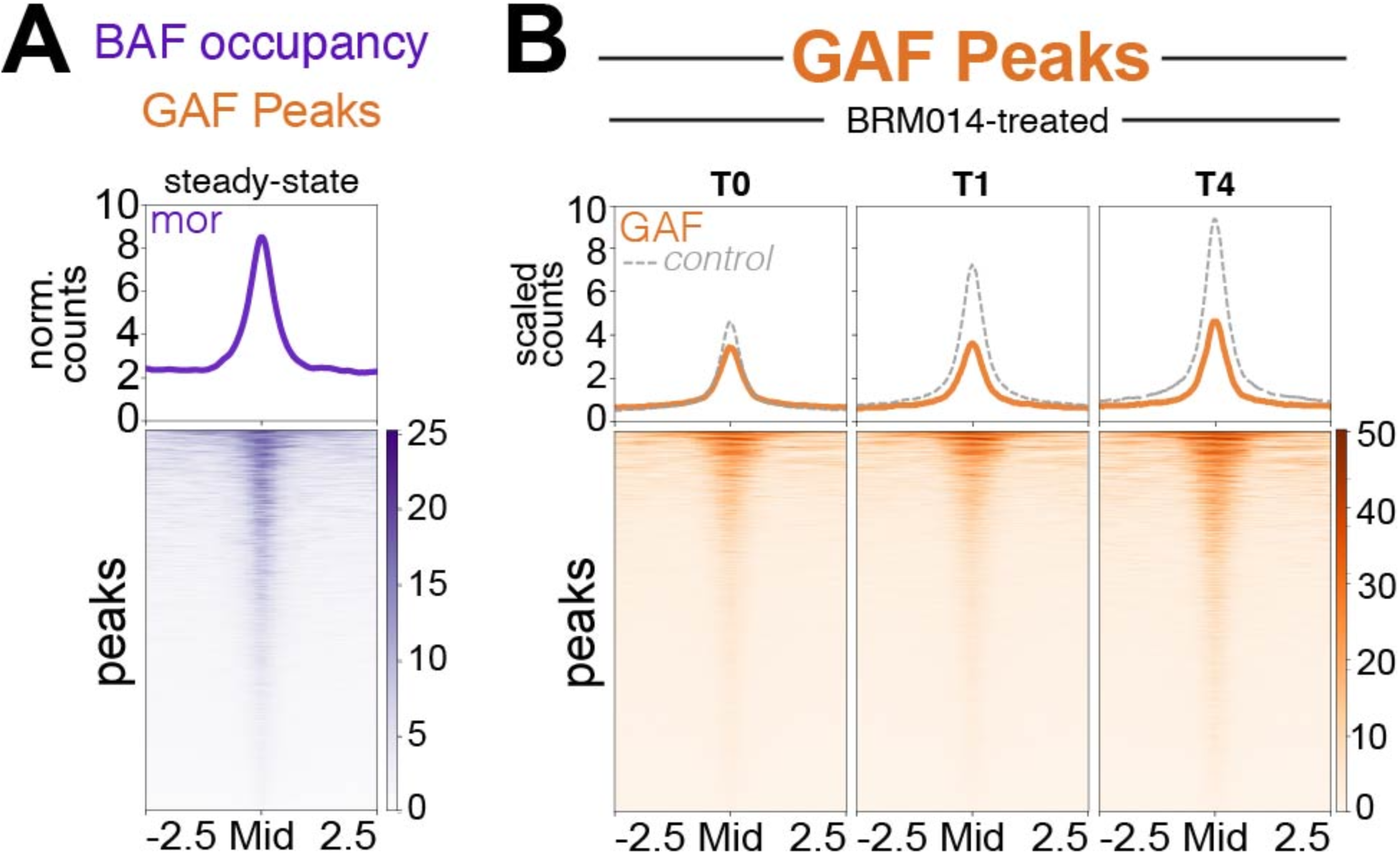
BAF inhibition disrupts GAF rebinding on nascent DNA. (**A**) Heatmap showing CUT&Tag profiling of Moira (BAF component) coverage at all GAF peaks. (**B**) Heatmap aligned to the center of all GAF peaks showing GAF signal over the course of Nascent CUT&Tag experiment treated with 10uM of BRM014. Grey dotted line indicates scaled counts of control data. Merged data of two biological replicates for Moira CUT&Tag, four biological replicates for BRM014-treated T0, two biological replicates for BRM014-treated T1 and two biological replicates for BRM014-treated T4.

To directly test if chromatin remodeling is required for GAF binding on nascent chromatin, we inhibited the BAF complex by treating Kc167 cells with BRM014 for 1 hour. BRM014 prevents ATP hydrolysis specifically by the Brahma (Brg1 homolog) catalytic subunit^21^. We then performed Nascent CUT&Tag profiling of GAF. Across all GAF sites on newly replicated chromatin, we observed only a subtle decrease in GAF binding immediately after replication fork passage, but larger decreases from parallel controls 1 hour and 4 hours after replication (**Figure 4B**). Thus, full restoration of GAF across the genome requires BAF activity.

### BAF regulates transcription factor occupancy on bulk chromatin

The loss of GAF binding observed during nascent chromatin maturation suggests that chromatin remodeling might be continually required for factor binding outside of S phase. To investigate how BAF inhibition impacts GAF binding on bulk chromatin, we treated an asynchronous population of Kc167 cells with BRM014 for one hour and profiled GAF binding with standard CUT&Tag. We observed that BRM014 treatment resulted in a ∼30% reduction of GAF binding genome-wide (**Figure 5A**), consistent with the idea that remodeling is continually required for stable GAF binding. Not all sites are equivalently affected: we identified 949 (13%) sites where GAF binding significantly decreases, indicating a strong dependence on chromatin remodeling (**Figure 5B,C**). Many of the features strongly affected by BRM014 treatment are developmental enhancers (47%), consistent with previous studies^22^ (**Figure 5D, Supplementary Figure 2**).

**Figure 5.**
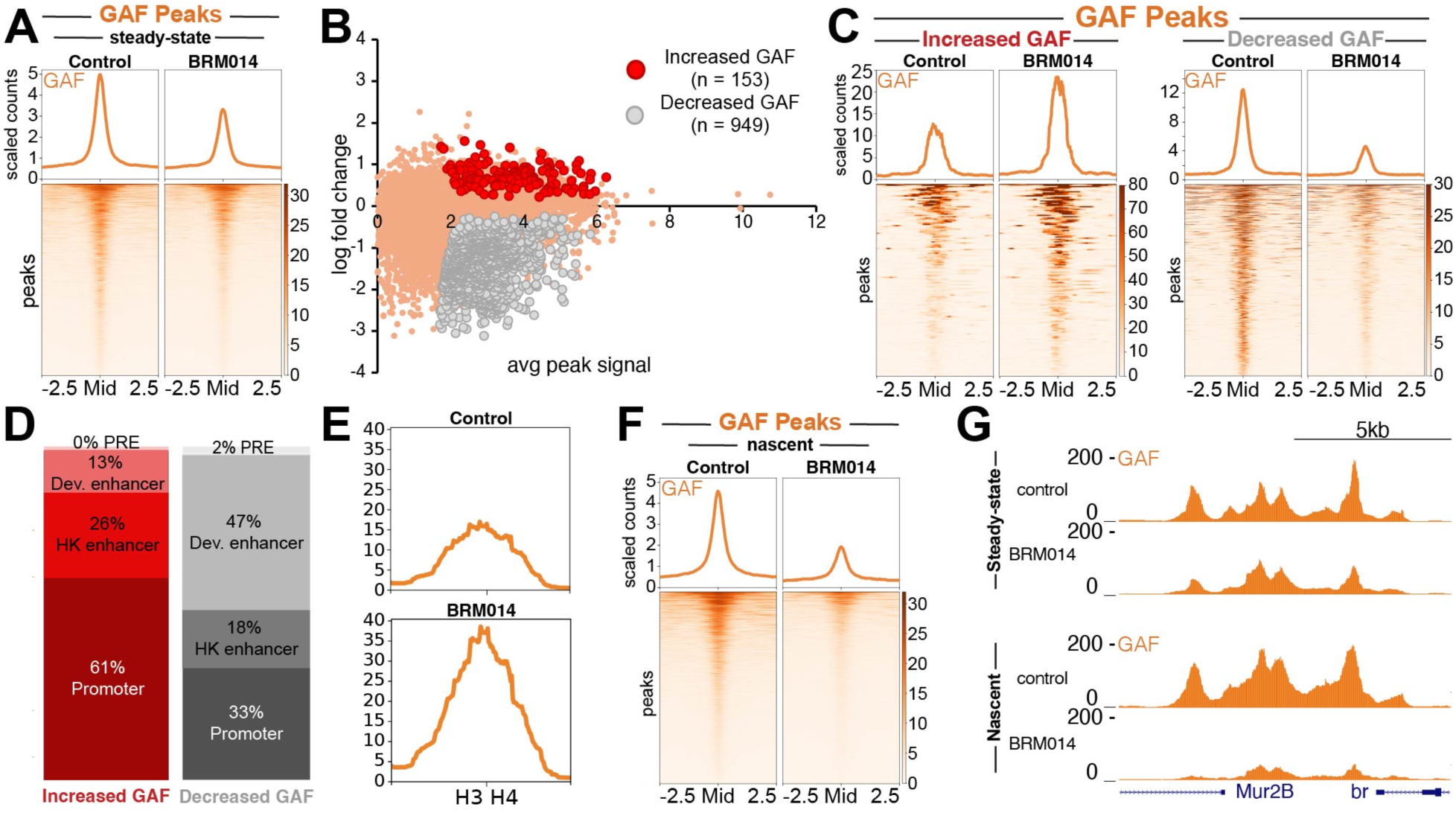
BAF regulates TF occupancy on nascent and bulk chromatin. (**A**) Heatmap aligned to the center of all GAF peaks showing CUT&Tag GAF signal in control and 1hr BRM014-treated samples. Read counts are scaled by number of mapped reads. (**B**) MA plot showing fold change in normalized counts comparing control to BRM014-treated samples. Statistical significance determined by two-sided t-test. (**C**) Heatmap aligned to the center of all GAF peaks gaining and losing GAF signal following BRM014-treatment showing GAF signal in control *versus* BRM014-treated samples. (**D**) Bar graph depicting proportion of features showing gains and losses in GAF signal following BRM014-treatment. (**E**) GAF coverage at bidirectional promoter element between histones H3 and H4. (**F**) Heatmap aligned to the center of all GAF peaks showing Nascent CUT&Tag GAF signal in control and 1hr BRM014-treated samples. Read counts are scaled by number of mapped reads. (**G**) UCSC browser track snapshot of CUT&Tag and Nascent CUT&Tag experiments centered on the *broad* promoter showing GAF signal for control and BRM014-treated samples.

Interestingly, we also observed 153 sites that show increased GAF signal upon BAF inhibition (**Figure 5B,C; Supplementary Figure 2**). The majority (61%) of these sites are gene promoters (**Figure 5D**), such as the H3-H4 bidirectional promoter element (**Figure 5E**, whose accessibility is normally maintained by chromatin remodelers other than BAF^22^.

In order to compare the effects of BAF inhibition on nascent *versus* bulk chromatin, we next used Nascent CUT&Tag to assess the impact of BAF inhibition on GAF binding to nascent chromatin after 1hr of BRM014 treatment. Globally, we observed that BAF inhibition results in a greater loss of GAF binding on nascent chromatin (**Figure 5F,G**) when compared to bulk chromatin. Given BAF’s nucleosome eviction activity, these data imply that TF binding sites become occluded by nucleosomes after replication fork passage and require BAF to evict nucleosomes to fully reestablish binding.

### Late recovering sites require BAF activity to reestablish GAF binding on nascent DNA

To better understand differences between bulk and nascent chromatin following BAF inhibition, we compared loss of GAF occupancy in BRM014-treated samples compared to controls. We identified 366 peaks which showed substantially greater loss of GAF binding in nascent BAF inhibition compared to bulk chromatin. On nascent chromatin, these peaks showed near total loss of GAF binding without BAF remodeling (**Figure 6A**), implying that these sites become nucleosome-occluded following replication fork passage, and that without BAF to evict nucleosomes, GAF is unable to reestablish stable binding. We therefore refer to these sites as nascent occluded sites. We next sought to understand how GAF binding at nascent occluded sites changes over the course of chromatin maturation with and without BAF inhibition. In control conditions, nascent-occluded sites were found to be late recovering peaks, with low levels of GAF occupancy immediately after replication fork passage, but substantial increases in GAF occupancy one hour (T1) and four hours (T4) after replication fork passage (**Figure 6B**).

**Figure 6.**
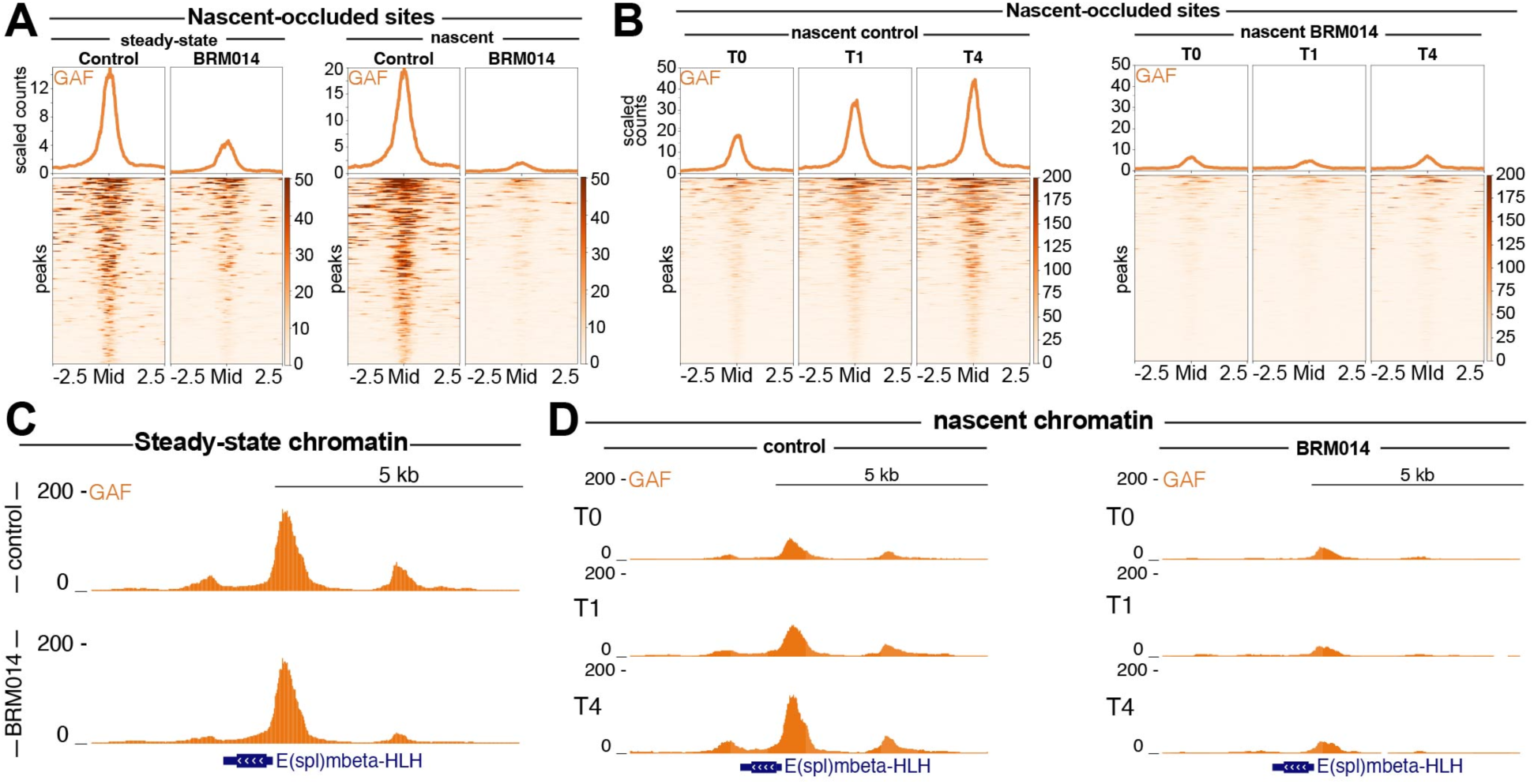
Nascent-occluded peaks are late recovering and highly sensitive to BAF inhibition. (**A**) Heatmaps of CUT&Tag and Nascent CUT&Tag data aligned to the center of nascent occluded peaks showing greater loss of GAF signal in Nascent CUT&Tag experiments compared bulk experiments. (**B**) Heatmaps of control and BRM014-treated Nascent CUT&Tag time course data aligned to the center of nascent occluded peaks. (**C**) UCSC browser snapshot of steady state CUT&Tag data showing control and BRM014-treated samples centered on the *Enhancer of split mβ* promoter (**D**) UCSC browser track snapshot centered on the *Enhancer of split mβ* promoter showing GAF signal across Nascent CUT&Tag time course.

Conversely, without BAF activity, these sites showed a substantial depletion in GAF occupancy relative to controls immediately after replication fork passage (T0), with no gains in GAF binding during the subsequent stages of chromatin maturation (T1, T4) (**Figure 6B**). These patterns can be observed at candidate peak regions, such as the promoter of the E(spl)mbeta-HLH (**Figure 6C-D**). Strikingly, while we saw little change in GAF occupancy at the E(spl)mbeta-HLH romoter in bulk conditions after BAF inhibition (**Figure 6C**), we observed substantial loss of GAF binding on nascent chromatin (**Figure 6D**), implying that at certain sites, BAF remodeling is critical for TF binding on nascent chromatin, but dispensable for sustained GAF occupancy on bulk chromatin. Together, these data suggest that nucleosomes deposited after the replication fork antagonize GAF binding, and that chromatin remodeling is essential to enable GAF binding to newly synthesized DNA.

### GAF binding increases at newly replicated GA-rich repeats

The 7,000 sites of GAF binding defined by peak calling include a subset of sites with extremely high signal in profiling (**Figure 7A**). These sites are centered in pericentric heterochromatin on stretches of AAGAG and other GA-rich repeat sequences, an abundant satellite repeat in the Drosophila genome^23^. These satellites have been shown to bind GAF^10^. As GAF coverage at these sites reflects multi-mapped reads across all GA-rich repeats in the genome, we summed these reads to assess GAF binding at satellite blocks. Surprisingly, Nascent CUT&Tag profiles reveal that GAF binding is higher immediately after replication fork passage and then decreases in later timepoints (**Figure 7B**). Newly replicated chromatin is transiently decondensed, which could provide a window of opportunity for GAF to bind before heterochromatin matures and compacts, thereby displacing GAF. Interestingly, GAF binding also increases substantially at GA-rich repeats following BAF inhibition (**Figure 7C**). As GAF is lost from many euchromatic sites during BRM014 treatment, GA-rich repeats might be acting as a sink for BAF freed from euchromatin.^24^

**Figure 7.**
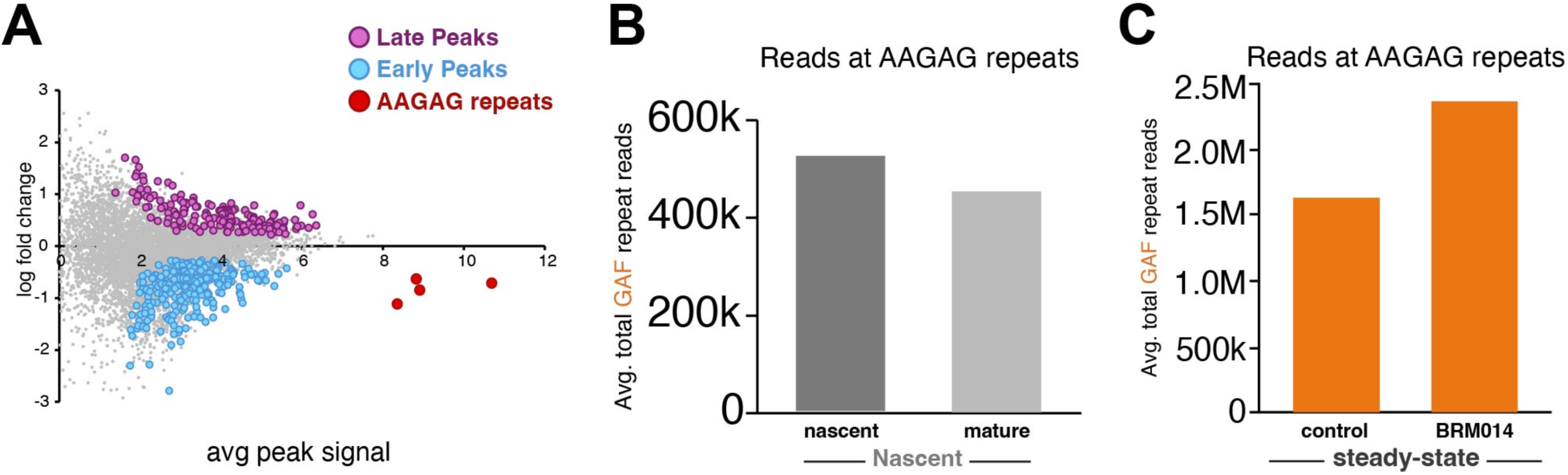
GAF binding increases at newly replicated GA-rich repeats and after BAF inhibition. (**A**) MA plot showing fold change in GAF occupancy T4 *versus* T0 with GA-rich repeats labelled in red (**B**) Total average reads at GA-rich repeats in nascent (T0) *versus* mature (T4) chromatin. (**C**) Total average reads at GA-rich repeats in control *versus* BRM014-treated samples. Merged data of four biological replicates.

## Discussion

Here we present Nascent CUT&Tag, a method for profiling chromatin factors on newly synthesized DNA and quantitatively tracking maturation over time. We demonstrate that Nascent CUT&Tag is effective at profiling a diverse array of chromatin features, including transcription factors such as GAF and PHO. As TF binding sites become occluded by nucleosomes following passage of the replication fork, nascent chromatin profiling provides a window into the kinetics and mechanisms by which TFs compete with nucleosomes to reestablish binding. In studying GAF and PHO, we find that TF rebinding to nascent chromatin is heterogeneous, with GAF showing a mixture of early and late recovery rates, while PHO largely shows delayed recovery across most binding sites genome-wide. We observed that at late recovering PHO sites, GAF reestablishes occupancy on nascent DNA prior to PHO, revealing a hierarchical order of transcription factor binding. Previous *in-vivo* profiling studies have shown that that PHO frequently co-binds to its target sites with other transcription factors, including GAF^34^. Furthermore, in-vitro studies using a chromatinized PRE template have shown that GAF binding is essential for PHO to binding to its target sites^35^. Together, these data suggest that delayed PHO recovery on newly synthesized DNA is due to a reliance on co-binding with other transcription factors such as GAF to achieve stable occupancy.

The rates of recovery of distinct chromatin features on nascent chromatin are highly variable, ranging from nearly instantaneous to requiring hours to become fully reestablished^16,17,25^. The deposition of nucleosomes occurs immediately after replication fork passage^26,27^. However, once established on nascent chromatin, it takes time for nucleosomes to fully wrap DNA^28,29^ and reestablish positioning^4^. This process is highly variable between organisms: yeast reestablish nucleosome positioning minutes after replication fork passage^30,31^, while in metazoans, reestablishment of nucleosome positioning^5^ and DNA accessibility^32^ requires over an hour.

Interestingly, this recovery is site specific, as house-keeping features show rapid recovery of nucleosome positioning, while sites related to developmental functions show substantial delays^5^. Studies that have assessed TF binding indirectly have shown that TFs likely play a critical role in reestablishing nucleosome positioning on nascent chromatin^33^. However, the timing and molecular mechanisms by which individual transcription factors regain occupancy on newly synthesized DNA have remained unclear.

The GAF transcription factor is essential for generating open chromatin in cell culture^36^ and early development^37,38^, and at many sites GAF binds to DNA prior to chromatin accessibility^38^. While GAF can bind to reconstituted nucleosomes^39,40^, it possesses little intrinsic chromatin remodeling activity^41^. Instead, GAF appears to recruit chromatin remodelers^42^ to expose factor binding sites and initiate transcription^41,43^. Using a chemical genetics approach to inhibit BAF activity, we demonstrate that BAF-mediated chromatin remodeling is essential for GAF to fully reestablish binding on nascent and bulk chromatin. Given the established role of BAF in moving and evicting nucleosomes, these data imply that at many sites throughout the genome, nucleosomes occupancy is refractory to GAF binding, and that prolonged BAF activity is required for GAF to achieve stable binding.

Using Nascent CUT&Tag to examine GAF to newly synthesized DNA, we observed a wide spectrum of recovery rates ranging from minutes to hours. Differences in recovery rate were associated with differences in GAF motif structure, with early recovery peaks showing short GA-rich motifs and late recovery sites showing longer, more degenerate motifs. Late recovery peaks also show a broader distribution of GAF binding than early recovering sites. As GAF multimerizes and preferentially binds motif arrays relative to single sites^9,40,44,45^, the broad distribution of GAF at late recovering sites may be due to GAF oligomers bound across extended arrays. Together with our data showing that BAF activity is essential for GAF to fully reestablish binding on nascent chromatin, we propose the following model: After replication fork passage, BAF remodels nucleosomes deposited on new DNA. At early recovery sites, only one or two nucleosomes must be remodeled for GAF to bind a motif. Conversely, at late recovery sites, more extensive remodeling is required to expose the multiple motifs needed for full GAF oligomer binding. As late recovery sites are associated with developmental features, delays in recovery stemming from the need for extended BAF remodeling could aid in regulating selective gene activity required during development. Interestingly, studies that have mutated degenerate TF motifs to stronger consensus motifs at developmental features have observed spurious gene activation and disruptions to normal development^46^, emphasizing the importance of motif structure in developmental gene regulation.

We observed that certain regions gain GAF binding upon BAF inhibition, including the GA-rich repeats found in pericentromeric heterochromatin, suggesting that these genomic regions could act as a sink for GAF binding following drug treatment^24^. We also observed increased GAF binding at GA-rich repeats immediately after replication fork passage, after which GAF binding decreases at GA-rich repeats as heterochromatin matures and condenses. This pattern is reminiscent of GAF binding at satellites in mitotic cells^18^, where GAF becomes enriched at GA-rich repeats during mitosis, and then redistributes to binding sites in euchromatin during chromatin decondensation in interphase cells. Together, these observations suggest that regardless of the mechanism that liberates GAF from DNA, GA-rich heterochromatic repeats have the potential to act as a sink for unbound GAF, perhaps limiting spurious binding events.

Alternatively, it is possible that GAF binding to newly replicated GA-rich repeats could help to reestablish heterochromatin structure disrupted by replication fork passage, as GAF binding is required to establish repressive heterochromatin at pericentromeric repeats during early *Drosophila* embryogenesis^10^. Interestingly, such a mechanism has also been proposed for the mammalian transcription factor Pax3^47^ which also binds a large repetitive satellite array in human cells, suggesting that TF binding to newly synthesized repeats may be a general mechanism for reestablishing heterochromatin disrupted by replication fork passage.

Transcription factor binding sites are prevalent in satellite repeats, and it has been proposed that reiterated arrangement of transcription factor binding sites within repeat sequences is an intrinsic mechanism for heterochromatin formation^48^. In such cases, Nascent CUT&Tag provides a general tool for investigating the timing and mechanisms governing TF binding during heterochromatin replication and chromatin maturation.

## Materials and methods

### Cell culture and drug treatment

Drosophila Kc167 cells (RRID:CVCL_Z834) were grown to log phase in HYQ-SFX Insect medium (Invitrogen) supplemented with 18 mM l-glutamine and harvested as previously described (43). All cell counts and measures of cell size were measured using the Vi-CELL XR Cell Viability Analyzer (www.beckman.com). BRM inhibitor (BRM014) was resuspended to 10 mM in dimethyl sulfoxide and frozen in aliquots. For cell treatments, compounds were added to a cell medium containing 2.0 × 10^6 cells per ml to a final concentration of 10 BRM014 and incubated at room temperature (RT) for times indicated in each experimental paradigm. For nascent timepoints, BRM014 was added immediately before EdU labeling, and remained in media for the duration of the EdU pulse. For chase timepoints, BRM014 was included in all media used to wash and resuspend cells, meaning that chases of 1hr and 4hrs reflect treatment times of BRM014 of 1hr and 4hrs respectively. For steady state experiments, compounds were added to a cell medium containing 2.0 × 10^6 Kc167 cells per ml to a final concentration of 10 µM BRM014 and incubated at room temperature (RT) for times indicated in each experimental paradigm. Aliquots of 500K cells were taken for bulk chromatin CUT&Tag profiling

### Nascent CUT&Tag

Kc167 were grown to confluence and split down to a concentration of one million Drosophila Kc167 cells per ml 20 hours before the start of EdU labelling. EdU analogue was then added to the media at a concentration of 10uM and incubated for 15 minutes. Cells were then spun down and resuspended in 30mL cell culture media containing 20uM cold thymidine. Cells were spun down again and resuspended in cell culture chase media (equal parts fresh media and conditioned media + 20uM thymidine). A volume was taken corresponding to 2 million cells per ml at current cell density and placed on ice as the nascent timepoint sample. The remainder of the cells were left to chase for timepoints specified. Additional volumes of samples were taken 1hr post EdU addition and 4 hours post EdU addition. Once all samples were taken, CUT&Tag with phenol-chloroform DNA extraction was performed as described^6,49–51^. Following extraction, DNA was solubilized in 10uL of H2O. Click chemistry was then performed to conjugate biotin onto replicated DNA molecules. Following completion of click-chemistry, SPRI bead cleanup was performed and DNA was resuspended in 20uls of water. Replicated DNA was then pulled down using Streptavidin T1 beads according to manufacturer’s instructions. Following completion of pull-down 21 µL of DNA bound to streptavidin T1 beads was mixed with a universal i5 and a uniquely barcoded i7 primer and amplified with NEB Q5 high-fidelity 2× master mix (catalog no. M0492S). The libraries were purified with 1.1× volume of Sera-Mag carboxylate-modified magnetic beads and subjected to LabChip DNA analysis and Illumina sequencing. Barcoded Nascent CUT&Tag libraries were pooled at equal volumes to normalize read counts between samples.

### CUT&Tag data processing and analysis

Libraries were sequenced on an Illumina NovaSeq instrument with paired-end 50 × 50 reads Barcoded CUTAC libraries were pooled at equal volumes within group to allow for normalization of total reads across samples.

Adapters were clipped by cutadapt (http://dx.doi.org/10.14806/ej.17.1.200) version 2.9 with the following parameters:

-j 8 --nextseq-trim 20 -m 20 -a AGATCGGAAGAGCACACGTCTGAACTCCAGTCA -A AGATCGGAAGAGCGTCGTGTAGGGAAAGAGTGT -Z. Clipped reads were aligned by Bowtie2^52^ to the UCSC D. melanogaster Dm6 reference sequence^53^ with the following parameters:

--very-sensitive-local --soft-clipped-unmapped-tlen --dovetail --no-mixed --no-discordant -q --phred33 -I 10 -X 1000. Clipped reads were also aligned by Bowtie2^52^ to the UCSC D. melanogaster Dm6 reference sequence^53^ with the following parameters: --end-to-end --very-sensitive --no-overlap --no-dovetail --no-mixed --no-discordant -q --phred33 -I 10 -X 1000. Properly paired reads were extracted from the alignments by SAMTools (version 1.9)^54^. Normalized count tracks in bigwig format were also made by bedtools^55^ 2.30.0 genomecov which are the fraction of counts at each base pair scaled by the size of the reference sequence (137,567,484) so that if, the scaled counts were uniformly distributed, there would be 1 at each position. Bigwig files were then uploaded to Galaxy^56^ and heatmaps were generated using the computematrix function in deepTools (version 3.5.1)^57^.

A bedfile containing a list of all D. melanogaster promoters, enhancers and PREs was used by computematrix function to compare scaled normalized reads from bedgraph files aligned at midpoint all D. melanogaster promoters, enahancers and PREs. The output of the computematrix function was visualized using the plotHeatmap function in Galaxy.

To generate plots showing changes in the GAF coverage in control versus BRM014-treated cells in nascent vs. steady state experiments, the total GAF signal at each peak promoter from 500-bp upstream to 500-bp downstream was summed using the multiBigwigSummary function from Galaxy deepTools.

### Generating scaling factors

To generate a scaling factor to normalize read counts between nascent timepoints, we divided the total mapped reads of each sample in the experiment by the average number of reads at the nascent timepoint. We generated scaling values for each timepoint across all experiments. We then averaged scaling values at each timepoint across experiments to generate the averaged scaling value for that timepoint. We used the averaged scaling value in the computeMatrix function to scale the normalized read counts for each timepoint accordingly.

To generate a scaling factor to normalize read counts between control and BRM014-treated samples, we divided the total mapped reads in each paired BRM014-treated and control sample and averaged the values. We used the averaged scaling value in the computeMatrix function to scale the normalized read counts for each timepoint accordingly.

### Identifying nascent sensitive peaks

We identified nascent sensitive peaks first by summing total signal at each peak in control and BRM014 treated conditions in steady state and nascent experiments. We then filtered our lists to contain only peaks that showed statistically significant differences between control and drug treated conditions using two-sided t-tests. We then calculated the absolute difference by subtracting the total signal in control conditions by BRM014-treated conditions for steady state and nascent samples. We then sorted peaks by those which showed losses greater than 5 arbitrary units in the nascent condition when compared to the steady state condition. Five was chosen, as this was the approximate inflection point in a knee plot of all differences between nascent and bulk chromatin. We also generated a fold change value by dividing the signal in control groups by signal in BRM014-treated groups. We then compared those ratios between nascent and bulk chromatin experiments. Peaks in which the fold change in nascent samples was 1.5-fold greater than in bulk chromatin experiments we deemed to be sensitive in nascent. The 1.5-fold threshold was selected as this was the approximate inflection point in a knee plot of nascent *versus* bulk chromatin ratios. Peaks identified by raw difference and fold change were merged and duplicates removed to generate the final nascent-sensitive list of peaks.

## Acknowledgments

We thank our Fred Hutchinson Cancer Center colleagues Doris Xu for help with cell culture, Christine Codomo and Terri Bryson for sequencing library pooling, and Jorja Henikoff for preparing the sequencing data for analysis.

## Author contributions

Conceptualization: M.W., K.A., and S.H. Methodology: M.W., K.A., and S.H. Investigation: M.W. K.N. and B.T. Visualization: M.W. Supervision: K.A. and S.H. Writing—original draft: M.W. Writing—review and editing: M.W., K.A., and S.H.

---

**Supplementary Figure 1.**
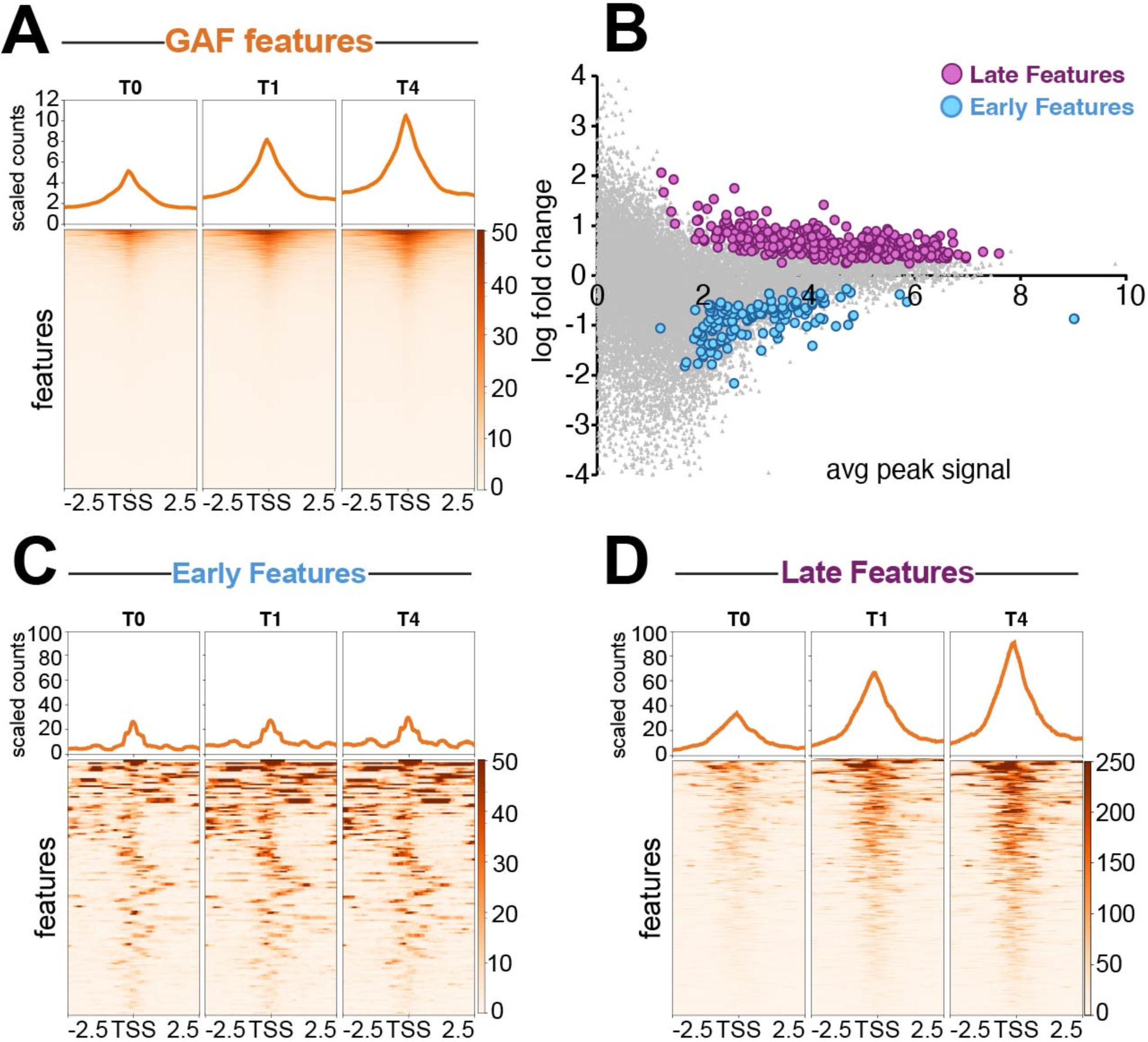
GAGA factor is lost on nascent chromatin and recovers over time. (**A**) Heatmap aligned to the center of all *Drosophila* features showing GAF signal over the course of Nascent CUT&Tag experiment. Read counts are scaled by number of mapped reads. (**B**) MA plot showing fold change in normalized counts comparing T4 to T0. Statistical significance determined by two-sided t-test. (**C**) Heatmap aligned to the center of all early recovering GAF features showing GAF signal over the course of Nascent CUT&Tag experiment (**D**) Heatmap aligned to the center of late recovering GAF features showing GAF signal over the course of Nascent CUT&Tag experiment.

**Supplementary Figure 2.**
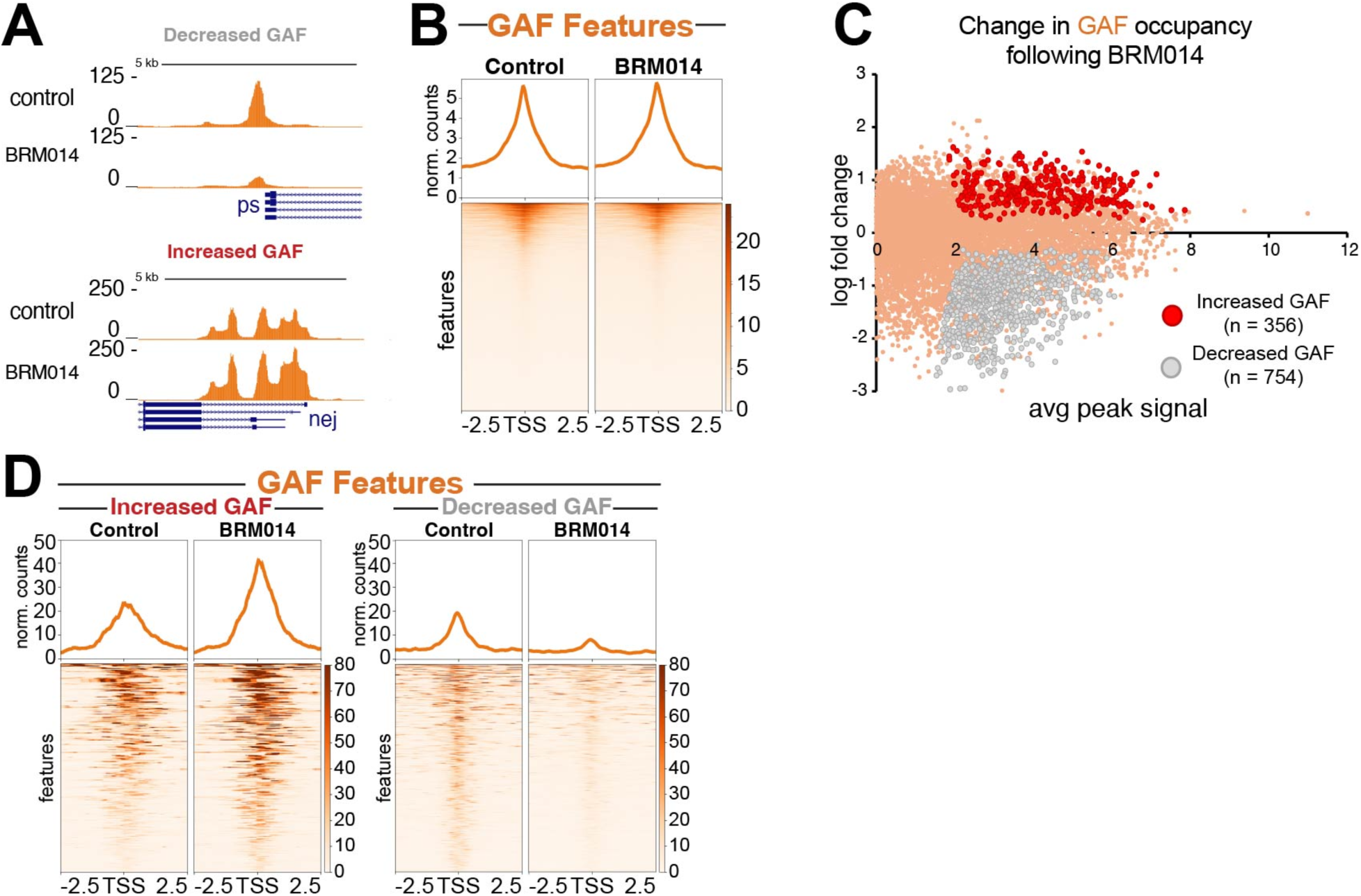
BAF inhibition alters GAF binding on bulk chromatin. (A) Representative UCSC browser track snapshot of control and BRM014-treated samples showing GAF signal at peaks gaining and losing GAF signal following BRM014-treatment. (B) Heatmap aligned to the center of all GAF features showing CUT&Tag GAF signal in control and 1hr BRM014-treated samples. Read counts are scaled by number of mapped reads (**C**) MA plot showing fold change in normalized counts at features comparing control to BRM014-treated samples. Statistical significance determined by two-sided t-test. (**D**) Heatmap aligned to the center of all GAF features gaining and losing GAF signal following BRM014-treatment showing GAF signal in control *versus* BRM014-treated samples.

## References

1 Luger, K., Dechassa, M. L. & Tremethick, D. J. New insights into nucleosome and chromatin structure: an ordered state or a disordered affair? Nat Rev Mol Cell Biol 13, 436–447 (2012). 10.1038/nrm3382

2 Maston, G. A., Evans, S. K. & Green, M. R. Transcriptional Regulatory Elements in the Human Genome. Annual Review of Genomics and Human Genetics 7, 29–59 (2006). 10.1146/annurev.genom.7.080505.115623

3 Spitz, F. & Furlong, E. E. M. Transcription factors: from enhancer binding to developmental control. Nature Reviews Genetics 13, 613–626 (2012). 10.1038/nrg3207

4 Ramachandran, S., Ahmad, K. & Henikoff, S. Capitalizing on disaster: Establishing chromatin specificity behind the replication fork. BioEssays 39, 1600150 (2017). 10.1002/bies.201600150

5 Ramachandran, S. & Henikoff, S. Transcriptional Regulators Compete with Nucleosomes Post-replication. Cell 165, 580–592 (2016). 10.1016/j.cell.2016.02.062

6 Kaya-Okur, H. S. et al. CUT&Tag for efficient epigenomic profiling of small samples and single cells. Nature Communications 10, 1930 (2019). 10.1038/s41467-019-09982-5

7 Farkas, G. et al. The Trithorax-like gene encodes the Drosophila GAGA factor. Nature 371, 806–808 (1994). 10.1038/371806a0

8 Mahmoudi, T., Katsani, K. R. & Verrijzer, C. P. GAGA can mediate enhancer function in trans by linking two separate DNA molecules. The EMBO journal 21, 1775–1781 (2002). 10.1093/emboj/21.7.1775

9 Chetverina, D., Erokhin, M. & Schedl, P. GAGA factor: a multifunctional pioneering chromatin protein. Cell Mol Life Sci 78, 4125–4141 (2021). 10.1007/s00018-021-03776-z

10 Gaskill, M. M. et al. Localization of the Drosophila pioneer factor GAF to subnuclear foci is driven by DNA binding and required to silence satellite repeat expression. Developmental cell 58, 1610–1624.e1618 (2023). 10.1016/j.devcel.2023.06.010

11 Brown, J. L., Mucci, D., Whiteley, M., Dirksen, M. L. & Kassis, J. A. The Drosophila Polycomb group gene pleiohomeotic encodes a DNA binding protein with homology to the transcription factor YY1. Mol Cell 1, 1057–1064 (1998). 10.1016/s1097-2765(00)80106-9

12 Brown, J. L., Price, J. D., Erokhin, M. & Kassis, J. A. Context-dependent role of Pho binding sites in Polycomb complex recruitment in Drosophila. Genetics 224 (2023). 10.1093/genetics/iyad096

13 Langlais, K. K., Brown, J. L. & Kassis, J. A. Polycomb Group Proteins Bind an engrailed PRE in Both the “ON” and “OFF” Transcriptional States of engrailed. PloS one 7, e48765 (2012). 10.1371/journal.pone.0048765

14 Kubota, T., Katou, Y., Nakato, R., Shirahige, K. & Donaldson, A. D. Replication-Coupled PCNA Unloading by the Elg1 Complex Occurs Genome-wide and Requires Okazaki Fragment Ligation. Cell Rep 12, 774–787 (2015). 10.1016/j.celrep.2015.06.066

15 Zhang, Z., Shibahara, K.-i. & Stillman, B. PCNA connects DNA replication to epigenetic inheritance in yeast. Nature 408, 221–225 (2000). 10.1038/35041601

16 Reverón-Gómez, N. et al. Accurate Recycling of Parental Histones Reproduces the Histone Modification Landscape during DNA Replication. Mol Cell 72, 239–249.e235 (2018). 10.1016/j.molcel.2018.08.010

17 Alabert, C. et al. Two distinct modes for propagation of histone PTMs across the cell cycle. Genes Dev 29, 585–590 (2015). 10.1101/gad.256354.114

18 Platero, J. S., Csink, A. K., Quintanilla, A. & Henikoff, S. Changes in chromosomal localization of heterochromatin-binding proteins during the cell cycle in Drosophila. J Cell Biol 140, 1297–1306 (1998). 10.1083/jcb.140.6.1297

19 Crosby, M. A. et al. The trithorax group gene moira encodes a brahma-associated putative chromatin-remodeling factor in Drosophila melanogaster. Mol Cell Biol 19, 1159–1170 (1999). 10.1128/mcb.19.2.1159

20 Janssens, D. H. et al. Automated CUT&Tag profiling of chromatin heterogeneity in mixed-lineage leukemia. Nature Genetics 53, 1586–1596 (2021). 10.1038/s41588-021-00941-9

21 Chambers, C. et al. SWI/SNF Blockade Disrupts PU.1-Directed Enhancer Programs in Normal Hematopoietic Cells and Acute Myeloid Leukemia. Cancer Res 83, 983–996 (2023). 10.1158/0008-5472.Can-22-2129

22 Hendy, O. et al. Developmental and housekeeping transcriptional programs in Drosophila require distinct chromatin remodelers. Molecular Cell 82, 3598–3612.e3597 (2022). 10.1016/j.molcel.2022.08.019

23 Lohe, A. R. & Brutlag, D. L. Identical satellite DNA sequences in sibling species of Drosophila. J Mol Biol 194, 161–170 (1987). 10.1016/0022-2836(87)90365-2

24 Spofford, J. B. Position-effect variegation in Drosophila. In The genetics and biology of Drosophila (ed. M. Ashburner and E. Novitski),. Academic Press, New York. vol. 1, pp. 955–1018. (1976.).

25 Annunziato, A. T. Assembling chromatin: the long and winding road. Biochim Biophys Acta 1819, 196–210 (2013). 10.1016/j.bbagrm.2011.07.005

26 McKnight, S. L. & Miller, O. L. Electron microscopic analysis of chromatin replication in the cellular blastoderm drosophila melanogaster embryo. Cell 12, 795–804 (1977). 10.1016/0092-8674(77)90278-1

27 Serra-Cardona, A. & Zhang, Z. Replication-Coupled Nucleosome Assembly in the Passage of Epigenetic Information and Cell Identity. Trends Biochem Sci 43, 136–148 (2018). 10.1016/j.tibs.2017.12.003

28 Ostrowski, M. S. et al. The single-molecule accessibility landscape of newly replicated mammalian chromatin. Cell 188, 237–252.e219 (2025). 10.1016/j.cell.2024.10.039

29 Annunziato, A. T. & Seale, R. L. Histone deacetylation is required for the maturation of newly replicated chromatin. J Biol Chem 258, 12675–12684 (1983).

30 Fennessy, R. T. & Owen-Hughes, T. Establishment of a promoter-based chromatin architecture on recently replicated DNA can accommodate variable inter-nucleosome spacing. Nucleic Acids Research 44, 7189–7203 (2016). 10.1093/nar/gkw331

31 Vasseur, P. et al. Dynamics of Nucleosome Positioning Maturation following Genomic Replication. Cell Rep 16, 2651–2665 (2016). 10.1016/j.celrep.2016.07.083

32 Stewart-Morgan, K. R., Reverón-Gómez, N. & Groth, A. Transcription Restart Establishes Chromatin Accessibility after DNA Replication. Mol Cell 75, 284–297.e286 (2019). 10.1016/j.molcel.2019.04.033

33 Gutiérrez, M. P., MacAlpine, H. K. & MacAlpine, D. M. Nascent chromatin occupancy profiling reveals locus- and factor-specific chromatin maturation dynamics behind the DNA replication fork. Genome Res 29, 1123–1133 (2019). 10.1101/gr.243386.118

34 Orsi, G. A. et al. High-resolution mapping defines the cooperative architecture of Polycomb response elements. Genome research 24, 809–820 (2014). 10.1101/gr.163642.113

35 Mahmoudi, T., Zuijderduijn, L. M., Mohd-Sarip, A. & Verrijzer, C. P. GAGA facilitates binding of Pleiohomeotic to a chromatinized Polycomb response element. Nucleic Acids Res 31, 4147–4156 (2003). 10.1093/nar/gkg479

36 Fuda, N. J. et al. GAGA Factor Maintains Nucleosome-Free Regions and Has a Role in RNA Polymerase II Recruitment to Promoters. PLoS genetics 11, e1005108 (2015). 10.1371/journal.pgen.1005108

37 Bhat, K. M. et al. The GAGA factor is required in the early Drosophila embryo not only for transcriptional regulation but also for nuclear division. Development 122, 1113–1124 (1996). 10.1242/dev.122.4.1113

38 Gaskill, M. M., Gibson, T. J., Larson, E. D. & Harrison, M. M. GAF is essential for zygotic genome activation and chromatin accessibility in the early Drosophila embryo. eLife 10, e66668 (2021). 10.7554/eLife.66668

39 Tsukiyama, T., Becker, P. B. & Wu, C. ATP-dependent nucleosome disruption at a heat-shock promoter mediated by binding of GAGA transcription factor. Nature 367, 525–532 (1994). 10.1038/367525a0

40 Feng, X. A. et al. GAGA zinc finger transcription factor searches chromatin by 1D–3D facilitated diffusion. Nature Structural & Molecular Biology (2025). 10.1038/s41594-025-01643-0

41 Judd, J., Duarte, F. M. & Lis, J. T. Pioneer-like factor GAF cooperates with PBAP (SWI/SNF) and NURF (ISWI) to regulate transcription. Genes Dev 35, 147–156 (2021). 10.1101/gad.341768.120

42 Nakayama, T., Shimojima, T. & Hirose, S. The PBAP remodeling complex is required for histone H3.3 replacement at chromatin boundaries and for boundary functions. Development 139, 4582–4590 (2012). 10.1242/dev.083246

43 Duarte, F. M. et al. Transcription factors GAF and HSF act at distinct regulatory steps to modulate stress-induced gene activation. Genes Dev 30, 1731–1746 (2016). 10.1101/gad.284430.116

44 van Steensel, B., Delrow, J. & Bussemaker, H. J. Genomewide analysis of *Drosophila* GAGA factor target genes reveals context-dependent DNA binding. Proceedings of the National Academy of Sciences 100, 2580–2585 (2003). doi:10.1073/pnas.0438000100

45 Katsani, K. R., Hajibagheri, M. A. & Verrijzer, C. P. Co-operative DNA binding by GAGA transcription factor requires the conserved BTB/POZ domain and reorganizes promoter topology. Embo j 18, 698–708 (1999). 10.1093/emboj/18.3.698

46 Lim, F. et al. Affinity-optimizing enhancer variants disrupt development. Nature 626, 151–159 (2024). 10.1038/s41586-023-06922-8

47 Wu, T. F. et al. PAX3 loads onto pericentromeric heterochromatin during S phase through PARP1. Anticancer Res 34, 4717–4722 (2014).

48 Bulut-Karslioglu, A. et al. A transcription factor–based mechanism for mouse heterochromatin formation. Nature Structural & Molecular Biology 19, 1023–1030 (2012). 10.1038/nsmb.2382

49 Henikoff, S., Henikoff, J. G., Kaya-Okur, H. S. & Ahmad, K. Efficient chromatin accessibility mapping in situ by nucleosome-tethered tagmentation. eLife 9, e63274 (2020). 10.7554/eLife.63274

50 Kaya-Okur, H. S., Janssens, D. H., Henikoff, J. G., Ahmad, K. & Henikoff, S. Efficient low-cost chromatin profiling with CUT&Tag. Nature Protocols 15, 3264–3283 (2020). 10.1038/s41596-020-0373-x

51 Wooten, M., Takushi, B., Ahmad, K. & Henikoff, S. Aclarubicin stimulates RNA polymerase II elongation at closely spaced divergent promoters. bioRxiv (2023). 10.1101/2023.01.09.523323

52 Langmead, B. & Salzberg, S. L. Fast gapped-read alignment with Bowtie 2. Nat Methods 9, 357–359 (2012). 10.1038/nmeth.1923

53 Nassar, L. R. et al. The UCSC Genome Browser database: 2023 update. Nucleic Acids Res 51, D1188–d1195 (2023). 10.1093/nar/gkac1072

54 Danecek, P. et al. Twelve years of SAMtools and BCFtools. GigaScience 10 (2021). 10.1093/gigascience/giab008

55 Quinlan, A. R. BEDTools: The Swiss-Army Tool for Genome Feature Analysis. Curr Protoc Bioinformatics 47, 11.12.11–34 (2014). 10.1002/0471250953.bi1112s47

56 The Galaxy platform for accessible, reproducible, and collaborative data analyses: 2024 update. Nucleic Acids Res 52, W83–w94 (2024). 10.1093/nar/gkae410

57 Ramírez, F. et al. deepTools2: a next generation web server for deep-sequencing data analysis. Nucleic Acids Research 44, W160–W165 (2016). 10.1093/nar/gkw257

